# 3D vascular quantitation with application to computational modeling: a pre-clinical light sheet microscopy, high resolution ultrasound, nano-computed tomography comparison study

**DOI:** 10.64898/2026.03.13.711685

**Authors:** Daibo Zhang, Stephanie E. Lindsey

## Abstract

It is increasingly necessary to both study biology in 3D and obtain quantitative measurements. Not all 3D-reconstructions are created equal, particularly when using the anatomical model as a basis for force calculations, i.e. computational modeling. Here, we compare 3D anatomical reconstructions from two emerging imaging modalities: 4D ultrasound (4DUS) and light sheet fluorescent microscopy (LSFM) against our previous nano-computed tomography (nanoCT) cohort data, using the tortuous highly intricate pharyngeal arch artery system of the chick embryo as a test bed. We highlight modality-specific morphological image acquisition discrepancies and their influence on subsequent computational fluid dynamics results. Overall, LSFM accurately captured quantitative volumetric measurements of small rapidly-changing vascular morphologies while 4DUS systematically inflated small tortuous vessels. Differences in image-based morphology changes led to significant changes in computationally-obtained force magnitudes and flow patterns linked to vessel angle and tortuosity. This validates LSFM as a comparative preclinical vascular quantitative imaging tool and suggests that 4DUS needs extensive 3D anatomical validation for non cardiac chamber vessels.

## Introduction

Small animal vascular quantitation is becoming an increasingly important tool in the study of cardiovascular physiology and disease etiology. As scientists explore new disease models and dynamics, there is an increased need to visualize, quantify, and model vascular morphologies. Traditionally, biologists have used histology to provide detailed sections of internal and microscopic biological structures, even reconstructing serial sections to visualize configurations in 3D. Despite advances, histological analysis remains qualitative and is often limited to two dimensions. 3D reconstructions of serial sections are subject to alignment errors, artificial deformations and have proven challenging for the complex, tortuous, vascular morphologies commonly seen in development and diseased morphologies^1–3^. 3D quantitation is necessary to understand a structure’s full depth and spatial complexity. It requires accurately aligned and calibrated volume images. In the the clinical world, computed tomography (CT) and magnetic resonance imaging (MRI) are used to understand anatomy and morphology in a 3D context for diagnostic and treatment purposes.

Due to the elevated cost and limited accessibility of micro-MRI, biological scientists have most readily adopted micro/-nanoCT for high resolution quantitative imaging of small animal models^4–8^. Micro/nano-CT images, acquired by rotating x-ray beams around a subject’s body, are the most widely used imaging modality to capture small animal quantitative morphological images in uniform voxel format. Through MicroCT, the user is able to generate rapid and inexpensive quantitative 3D datasets with the help of exogenous contrast media. Although live CT imaging is possible^9,10^, *ex vivo* quantitative imaging techniques on preserved animal and tissue structures are more commonly used^5–8^. The resolution and/or duration of the live-imaging microCT window often do not offer an advantage over *ex vivo* micro/nanoCT imaging.

Microscopy and light sheet microscopy in particular^11–13^ has become common to biological imaging cores and is making quantitative imaging of small animal models faster and more accessible. Microscopy techniques often necessitate clearing to reduce light scatter for full network visualization^14,15^. Simple and inexpensive clearing methods such as iDISCO (immunolabeling-enabled three-dimensional imaging of solvent-cleared organs)^16^ allow for whole-mount immunolabeling with volume imaging of large cleared samples ranging from whole embryos to adult organs. iDISCO is modeled on classical histology techniques, facilitating translation of section staining assays to intact tissues. The original method was shown to be harsh on biological tissues with a tissue shrinkage of 50% when compared to MRI^12^. Thus iDISCO+ was developed to minimize tissue shrinkage via a milder dehydration procedure. A light sheet fluorescent microscopy (LSFM) iDISCO+ method was recently used to acquire quantitative high resolution images of the coronary vascular system in neonatal and adult mice for computational simulation^17^. We modified this technique, developing a rapid endo-DISCO light sheet preparation protocol for small animal flow quantitation^18^.

Advanced high-frequency small animal ultrasound is also becoming an increasingly popular trend for live quantitative imaging^19–24^, particularly for capturing wall motion and cardiac function. High-frequency ultrasound uses megahertz-frequency ultrasonic waves that scatter when transmitted or reflected through tissue. The ultrasound beam can be translated across planes to generate 3D images. When a 2D cine (time-loop) is captured at each plane, a 4 dimensional ultrasound (4DUS) data set (3-dimensional + time) is compiled. The cine data is particularly advantageous for gating and cardiac function measurements. From these high resolution images, 3D anatomical models can be created. While 4DUS has been shown to be a reliable and cost-effective technique for longitudinal studies of cardiac function and disease progression^25^, 3D vessel comparisons are lacking. The question remains, whether 3D reconstructions from 4DUS produce comparative resolution to CT images, when not being used to capture relative changes between experimental groups.

Quantitative analysis of 3D datasets generated by 4DUS, LSFM and nano/microCT techniques have not been compared. In this study, we compare 3D anatomical reconstructions from the two emerging imaging modalities, 4DUS and LSFM, against our previously published nano-computed tomography (nanoCT) dataset. Using the highly intricate pharyngeal arch artery (PAA) system of the chick embryo as a test-bed, we quantify how each imaging modality captures the rapid changes in cross-sectional area, and intricate morphology of the PAA system as a whole (Fig. 1). We subsequently perform subject-specific computational fluid dynamics modeling on each of modality captured cohort models to highlight the effect that morphology has on calculating mechanobiological forces such as pressure and wall shear stress.

**Figure 1.**
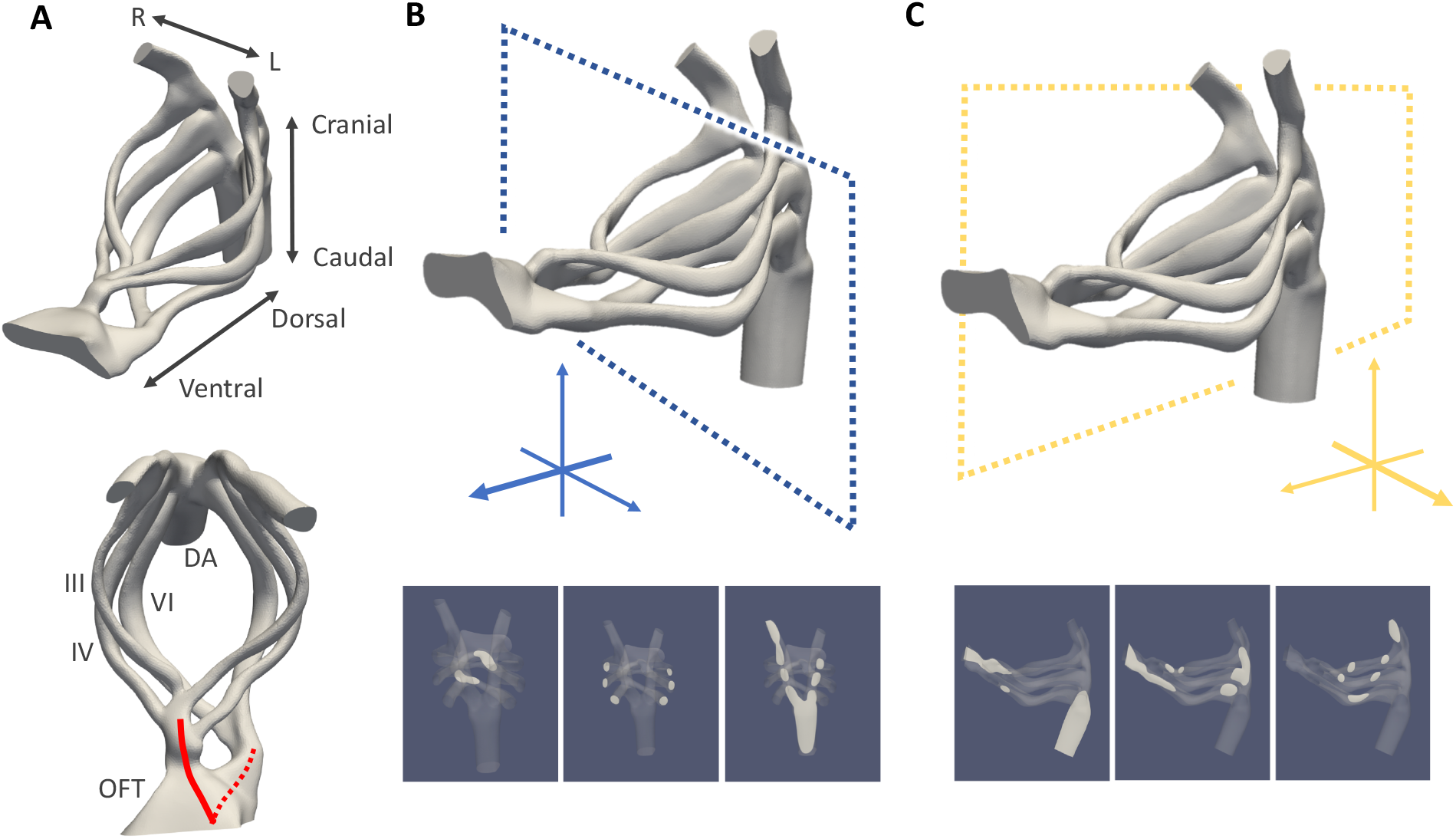
Anatomical landmark and imaging orientation guide. (A) Summary of the HH26 pharyngeal arch artery structure including its orientation within the body, position of the arches, and the partially septated OFT. (B) Schematics showing the imaging planes, normal to the ventral-dorsal axis, used for 4D ultrasound acquisition. Plane sections along arches shown below. (C) Schematics showing the imaging planes, normal to the left-right axis, used for light sheet acquisition. Plane sections along arches shown below.

## Results

### Quantitative imaging of the chick developing arch arteries

We performed quantitative imaging on HH26 (day 5) chick embryos’ arch artery system using LSFM and 4DUS and compared them against our team’s previous nanoCT imaging data^26^. The arch arteries were chosen because of their complex, tortuous geometry that rapidly changes in height and curvature (Figure 1). The imaging modalities of 4DUS and LSFM were chosen to determine whether either could serve as a more accessible alternative to nanoCT. To isolate the impact of imaging modality alone on 3D vascular morphology and subsequent subject-specific hemodynamic simulation results, we elected to image the PAAs from the same 5 embryos with both 4DUS and LSFM.

Sample prep, image acquisition and post-processing varied for each of the imaging modalities (Table 1, 2), with 4DUS capturing *in vivo* while LSFM and nanoCT captured perfusion-fixed embryos. 4DUS required the least sample preparation effort, but most significant active labor during image acquisition, as translation through the z-plane was not automated. As of date, simultaneous recording of B-mode cine loop and ECG in the chick embryo was technically impossible. Cardiac gating, motion correction, and plane-to-plane synchronization was performed retroactively based on ‘Spatial and Temporal Correlation,’ similar to^20^. We prepared embryos for LSFM imaging following a novel protocol^18^ that combines nonspecific labeling of vessel endothelium with FITC and tissue clearing through modified iDISCO+. LSFM and nanoCT sample preparations both demanded multiple days of sample preparation but had largely automated image acquisition procedures.

**Table 1.** Modality-based Tasks Required for High Resolution Image Acquisition.

| Modality | Sample preparation | Image acquisition | Post-processing |
| --- | --- | --- | --- |
| 4DUS | *Ex ovo* open culture | Manual probe translation<br>In-plane acquisition<br>(Repeat manually) | Retrospective gating<br>Synchronization<br>Intensity enhancement |
| LSFM | Sample staining<br>Sample fixation<br>Sample dehydration<br>Sample clearing | Sample mounting<br>In-plane acquisition<br>(Repeat automatically) |  |
| nanoCT | Sample fixation<br>Sample staining<br>Sample dehydration | Voxel acquisition |  |

**Table 2.** Comparison of the three imaging modalities in terms of capability and cost.

|  | **nanoCT** | **4DUS** | **LSFM** |
| --- | --- | --- | --- |
| Live imaging capability | Not possible | Widely capable | Limited to early chick or zebrafish embryos |
| Resolution (in-plane) | 0.15-25 $\mu\text{m}$ | 5-100 $\mu\text{m}$ | 0.1-5 $\mu\text{m}$ |
| Resolution (through-plane) | 0.15-25 $\mu\text{m}$<br>(= in-plane) | >20 $\mu\text{m}$<br>(≠ in-plane) | 2-4 $\mu\text{m}$<br>(≠ in-plane) |
| Max. depth of field | 80 mm | 15 mm | 5 mm<br>(without tiling) |
| Specialized skill | Embryo dissection, perfusion | Computer programming | Embryo dissection, perfusion |
| Sample prep time (per embryo active labor) | 15-45 min | 5-10 min | 30-60 min |
| Sample prep time (incubation) | 1-2 days | N/A | 2-3 days |
| Instrument user fee | \$200-400/hr | \$30-60/hr | \$20-36/hr |
| Imaging time (per embryo active labor) | <5 min | Approx. 30 min | Approx 10 min |
| Imaging time (per embryo instrument time) | Approx. 2 hr<br>(2-4 $\mu\text{m}$ resolution) | > 15 min | < 3 min<br>(1-3 $\mu\text{m}$ resolution) |
| Imaging stack file size | 1-3 GBs | 1-2 GBs (raw)<br><10 MBs (processed) | 3-5 GBs |

All imaging modalities were able to distinguish the PAAs from their surrounding tissues (Fig. 2), with NanoCT and LSFM images displaying stained lumen walls and/or surrounding tissue with dark lumen regions and 4DUS registering bright vessel lumens. In-plane LSFM images provide comparable sharpness and fine local details as nanoCT image, whereas 4DUS images contain noticeable noise within in-plane and through-plane views. The disparities between in-plane and through-plane resolutions of 4DUS and, to a lesser extent LSFM, introduced slight smudging/blurring artifacts in the final image stacks that did not interfere with 3D segmentation.

**Figure 2.**
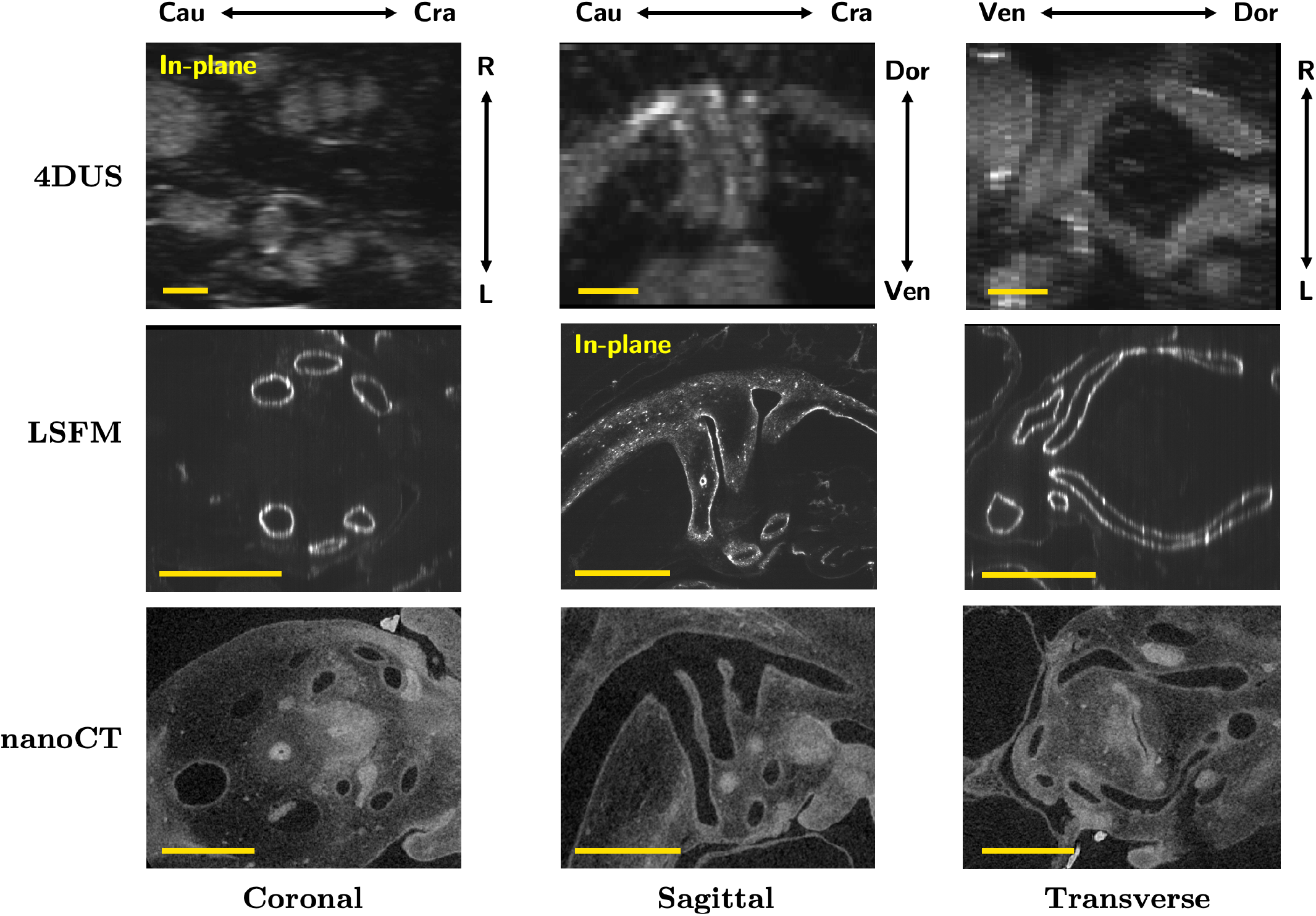
High resolution PAA image stacks. Post-processed images of the PAAs viewed through all three anatomical planes acquired by each imaging modality. In-plane image slices are indicated. Scale bars represent 0.5 mm. LSFM: light sheet fluorescence microscopy, 4DUS: four-dimensional ultrasound, nanoCT: nano computed tomography, Cau: caudal, Cra: cranial, R: right, L: left, Dor: dorsal, Ven: ventral.

### Morphology of 3D Reconstructions Observed Across Imaging Modalities

Following high resolution image acquisition, we used our ‘high-resolution imaging and flow measurement to multiscale numerical analysis pipeline’^26^ to obtain detailed morphological and hemodynamic comparisons between imaging modalities (Fig. 3). With this pipeline, 1) Subject-specific 3D anatomical models (*n* = 5) were reconstructed from high-resolution imaging stacks obtained from each imaging modality (Supplemental Figure S1). For 4DUS and LSFM, the same embryo obtained two scans, one for each modality, and therefore underwent two separate 3D anatomical reconstructions 2) Advanced morphometric analysis was performed along each of the six PAA vessels, obtaining morphological values (diameters, cross-sectional area, and ellipticity) at eleven point equally spaced points along each arch and 3) computational fluid dynamic modeling was performed.

**Figure 3.**
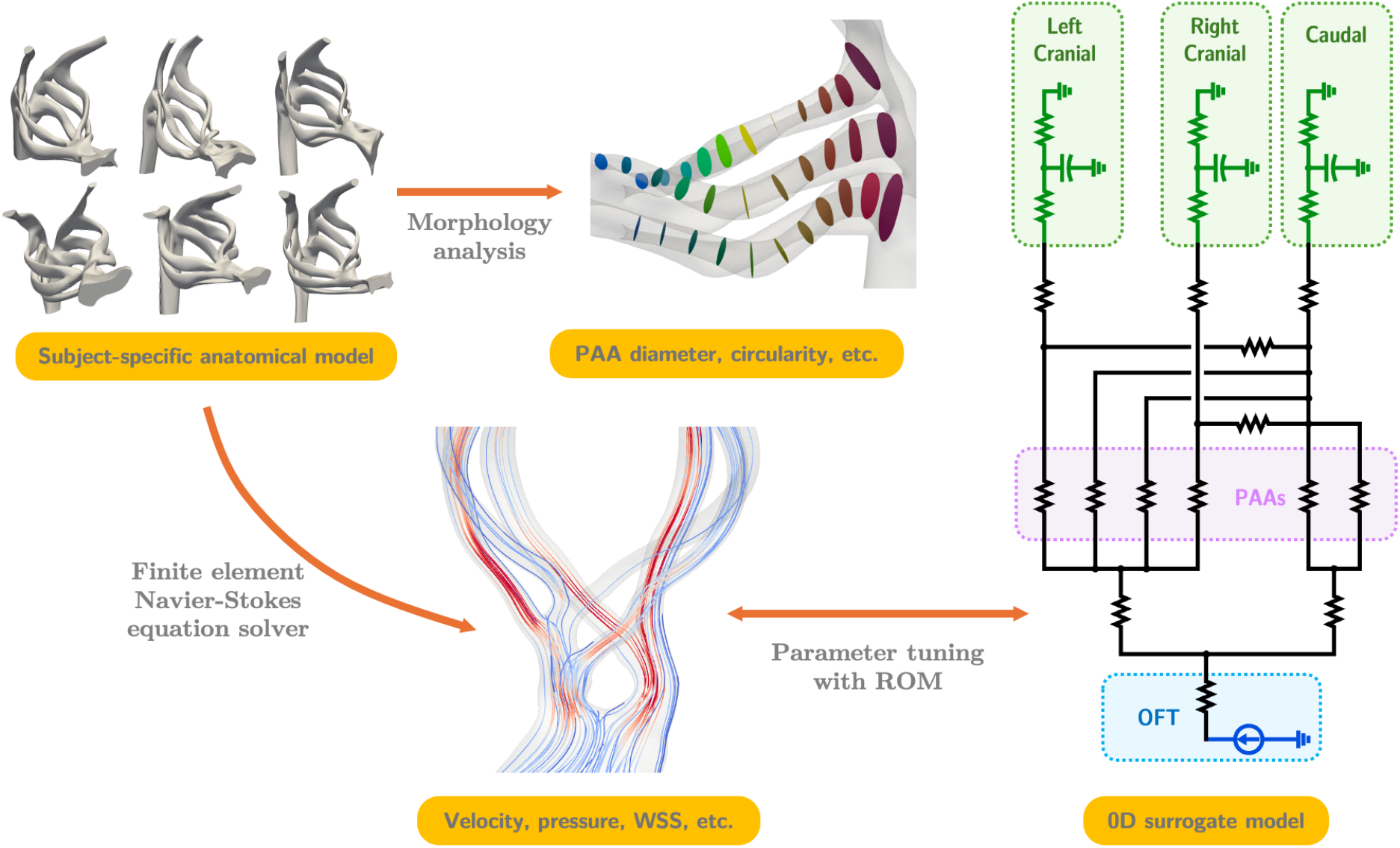
Workflow for anatomical model generation & morphological-hemodynamics multiscale analysis. Subject-specific anatomical reconstructions of the region of interest undergo detailed morphological analysis. The models are subsequently used to for pulsatile 3D hemodynamic simulations. A 0D surrogate model is used to tune the boundary conditions in the 3D model.

HH26 pharyngeal arch arteries are marked by lengthening, rotation and septation of the OFT and arch pairs^27,28^. All three imaging modalities captured the same overarching morphology: a septated OFT with upper aortic branches (PAAs III and IV) and caudal pulmonary trunks (PAAs VI). In the span of 24 hours of chick development, from HH24 (day 4) to HH26 (day 5), the OFT inlet has rotated from an in-line vertical position to a more lateral left and right position as seen in the representative models of Figure 4 (A). All three modalities recover similar branching angles for the aortic trunk and pulmonary trunk from the common OFT. While no statistically significant OFT rotation angles were found between groups, modality-based differences were observed between 4DUS and LSFM, where mean OFT rotation angles were found to be 66º and 82º respectively. Cohort-based differences in LSFM/4DUS and nanoCT rotation angles were also observed between groups, with 4DUS/LSFM OFT rotations consistently more advanced in the rotation cycle. The LSFM cohort-based mean OFT rotation angle of 82º, confirms that the aortic trunk has shifted almost exactly to the right of the pulmonary trunk, compared to nanoCT’s 35º angle. Cohort-based differences suggest that the 4DUS and LSFM embryos possess PAA structures in a slightly more advanced configuration within the HH26 stage. Despite the advanced rotation configuration, 4DUS/LSFM embryos were smaller in size than nanoCT cohort embryos. Such size effects are not uncommon even within the same cohort of embryos^20^. To facilitate relative arch comparisons, morphological values were normalized to arch length. Figure 4(B) shows the normalized lumen volume of each PAA.

**Figure 4.**
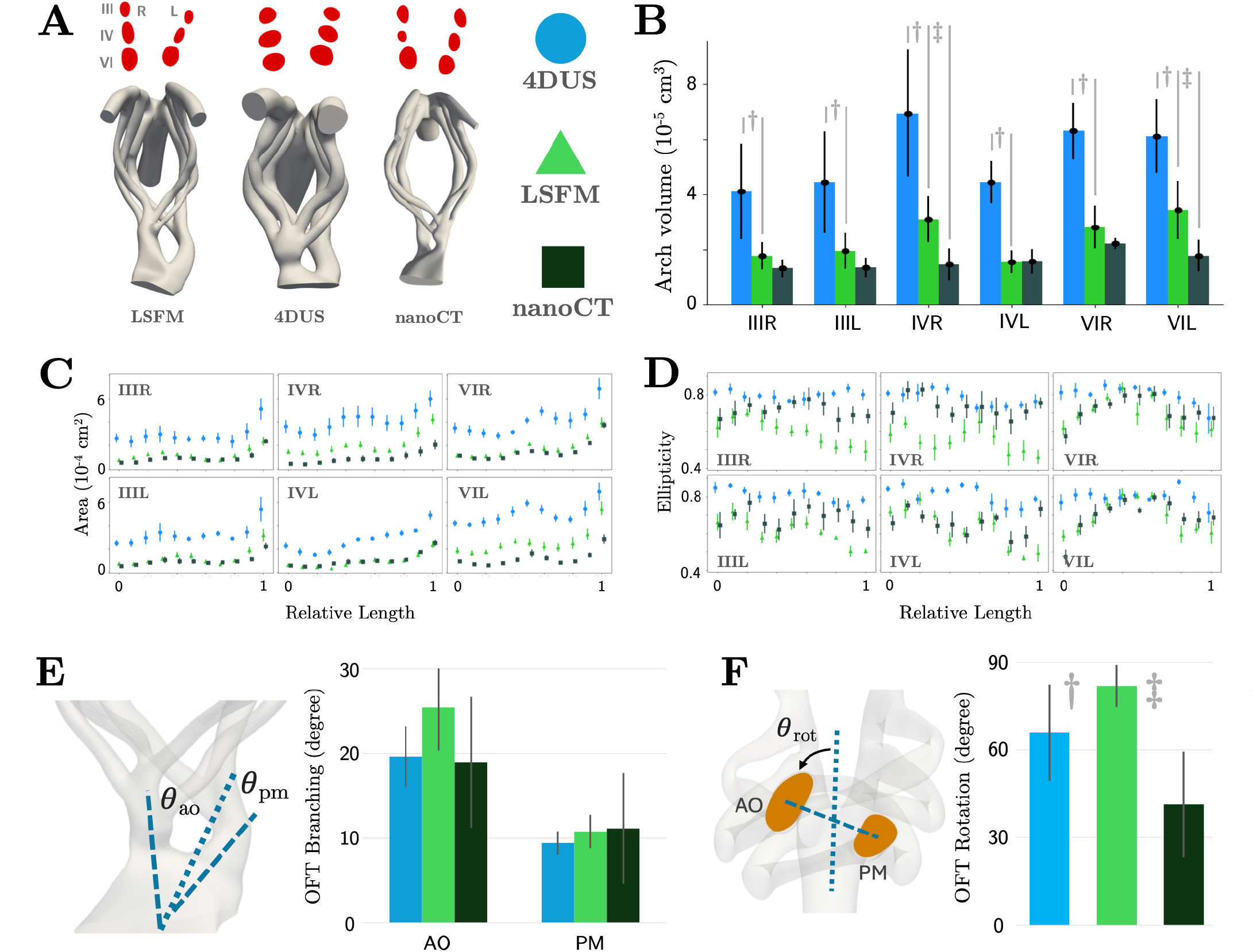
Control HH26 PAA mophology observed from the three imaging modalities. (A) Representative cross-sectional areas of the later half of the PAAs towards the dorsal root by modality and their representative 3D anatomical model. (B) PAA lumen volume measured from individual models constructed with each imaging modality, scaled by PAA length ratio across modalities. (C-D) cross-section (C) area (scaled based on arch length) and (D) ellipticity measured at various points along the PAAs from individual models of each modality. (E) OFT branching angle, the angle between the common OFT and the septated OFT branches (F) Angle of OFT rotation, the tilt of the AO-PM axis from the cranial-caudal axis of the embryo. LSFM: light sheet fluorescence microscopy, 4DUS: four-dimensional ultrasound, nanoCT: nano computed tomography, PAA: pharyngeal arch arteries, OFT: outflow tract, AO: aortic trunk, PM: pulmonary trunk. Data in (B,E,F) are presented as mean ± standard deviation, and *p*-values are indicated where statistical significances are found (*p* < 0.05, paired *t*-test for LSFM vs. 4DUS, Welch’s *t*-test for LSFM vs. nanoCT). Data in (C,D) are presented as mean ± standard error. See supplemental Figures S4-5 for individual replicate embryo data and statistical analysis.

The lower pixel resolution and higher noise level of 4DUS recordings tend to make the vessel appear thicker and rounder. As a result, 4DUS-derived volumes of all 6 arches are significantly larger than LSFM-derived volumes. 4DUS also displays noticeably larger CSAs and consistently more circular arches (Table 3, Supplemental figure S4-5), with an average ellipticity value above 0.8 (where 1 indicates a perfect circle) along the arches for all arch arteries.

**Table 3.** Number of sections (out of 11 per arch) showing statistical significant differences in each normalized morphological or hemodynamic metric of interest. *n* = 5 for all modalities. Statistical differences (*p* < 0.05) are assessed using paired *t*-test (4DUS vs. LSFM) and Welch’s *t*-test (LSFM vs. nanoCT). Locations of these sections and corresponding *p*-values are included in Supplemental figures S4-7.

|  | 4DUS vs. LSFM |  |  |  | LSFM vs. nanoCT |  |  |  |
| --- | --- | --- | --- | --- | --- | --- | --- | --- |
|  | CSA | Ellipticity | Pressure | WSS | CSA | Ellipticity | Pressure | WSS |
| IIIR | 5 | 8 | 7 | 1 | 0 | 3 | 0 | 2 |
| IIIL | 10 | 10 | 8 | 3 | 0 | 0 | 0 | 1 |
| IVR | 9 | 7 | 9 | 1 | 6 | 4 | 0 | 0 |
| IVL | 11 | 8 | 9 | 1 | 0 | 1 | 0 | 0 |
| VIR | 11 | 2 | 10 | 3 | 1 | 0 | 0 | 0 |
| VIL | 10 | 6 | 9 | 1 | 8 | 1 | 0 | 1 |

The LSFM models displayed prominently elliptical cross-sections elongating along the cranial-caudal axis, in overall agreement with nanoCT reconstructions. Some deviations between LSFM and nanoCT normalized arch volumes are seen in PAAs IVR and VIL (Fig. 4B). Light-sheet and nanoCT relative cross-sectional areas (CSAs) taken along the length of the arch largely follow similar trends with differences seen between PAA VIL and PAA IVR (Fig. 4C, Table 3, Supplemental figure S4). Despite differences in overall embryo size, LSFM consistently images the PAAs with the same shape and aspect ratio integrity as nanoCT. The more elliptical CSAs of LSFM embryos, compared to nanoCT, is in line with the more mature PAA configurations of LSFM embryos as determined by the more advanced OFT rotation. The increases in LSFM-derived PAA IVR and PAA VIL volume and CSAs are likely results of the OFT branch position differences.

### Resulting Modality-specific Computational Fluid Dynamics

Following our advanced morphometric analysis, we sought to characterize the extent to which imaging-associated anatomical disparities affect the quantification of flow and hemodynamic forces. The maximum Reynolds number associated with HH26 PAA blood flow is found to be around 30 (data not shown), which is lower than that of human fetal and adult arterial flow^29^. Nonetheless, pulsatile blood flow profiles within each PAA branch is expected to have a nontrivial relationship with the tortuous geometry of the PAA branch and its highly variable cross-sectional shapes^30^. In addition to cross-sectional area and perceived vessel resistance, flow distribution across the six arches depends on OFT orientation^31^. As such, values of key hemodynamic metrics cannot be directly inferred from morphology, necessitating the use of subject-specific blood flow simulations.

We performed subject-specific 3D pulsatile blood flow simulations in each PAA model. Cohort-based Doppler derived flow curves were used for the nanoCT and 4DUS/LSFM cohort models. Each simulation was tuned to achieve a 90/10 caudal/cranial flow split and measured caudal dorsal aortic pressure^26^. The resulting peak flow pressure and WSS distributions are shown in Figure 5(A) and Supplemental figure S3.

**Figure 5.**
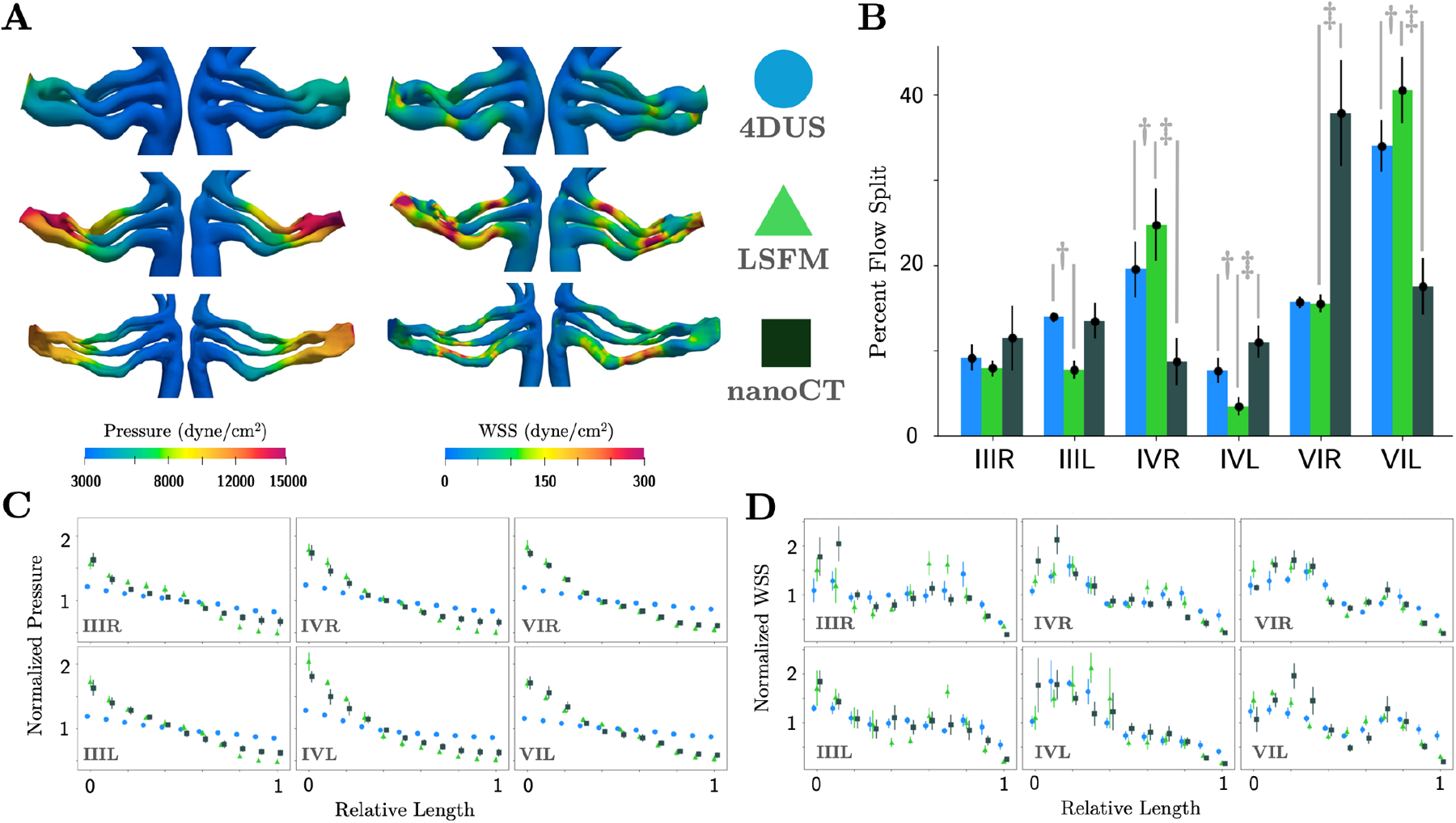
CFD hemodynamic simulation results of each imaging modality. (A) Peak flow pressure and WSS maps calculated from subject-specific simulation results in one representative PAA geometry for each modality. The 4DUS and LSFM models are of the same embryo. (B) Flow split across the six PAA branches for each imaging modality. Data presented as population mean ± standard deviation (*n* = 5). (C-D) Cross-sectional average time-maximum pressure (C) and WSS (D) along each arch, normalized to the average values of each metric along the corresponding arch. LSFM: light sheet fluorescence microscopy, 4DUS: four-dimensional ultrasound, nanoCT: nano computed tomography, WSS: wall shear stress. Data in (B) are presented as mean ± standard deviation, and *p*-values are indicated where statistical significances are found (*p* < 0.05, paired *t*-test for LSFM vs. 4DUS, Welch’s *t*-test for LSFM vs. nanoCT). Data in (C,D) are presented as mean ± standard error. See supplemental figures S6-7 for individual replicate embryo data and statistical analysis.

Flow split differed across all three imaging modalities (Fig. 5B). Although the same 4DUS/LSFM embryos were used, image-based flow simulations for 4DUS and LSFM geometries featured significantly different flow splits to PAAs IIIL, IVR, IVL, and VIL. When compared to LSFM, nanoCT-based simulations featured lower flow split to PAAs IVR and VIL and high flow split to PAAs IVL and VIR. Flow split differences likely resulted from differences in OFT angles^31^. Differences between nanoCT and the LSFM-4DUS embryos were expected due to the fact that the pulmonary trunk is shifted further to the left of the aortic branches in all LSFM-4DUS embryos (Fig. 4, Fig. S3). This alignment likely increases flow split to PAA IVR and VIL, subsequently reducing flow to IVL and VIR.

In all cases, pressure dissipates through the arch arteries before reaching the DoA. High pressure magnitudes travel further along LSFM arch lengths before reaching the stage specific lower range of 3700 dynes/cm^2^. Due to larger 4DUS CSAs, 4DUS arch artery systems displayed noticeably lower pressure and wall shear stress (WSS) magnitude values (Table 4, 5). Longer arches and larger overall PAA size of nanoCT embryos resulted in lower pressure and WSS magnitude compared to LSFM embryos (Fig. 5A, Fig. S3).

**Table 4.**
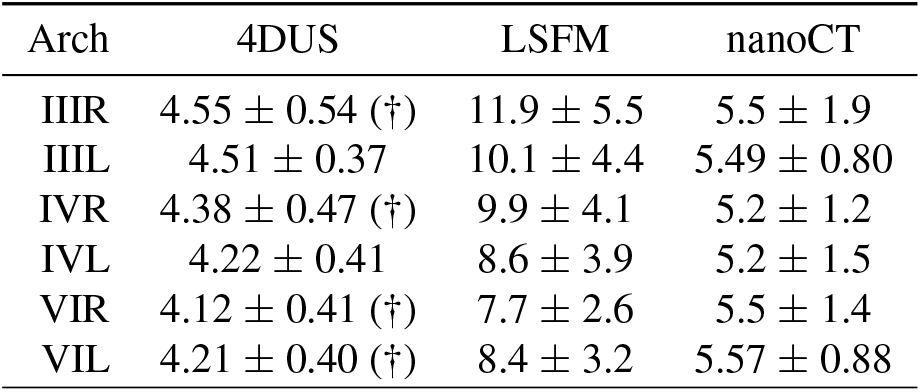
Raw values of spatial-mean peak systolic pressure (in 10^3^ dyne/cm^2^) across each arch based on each imaging modality. All data are presented as population mean ± standard deviation (*n* = 5). † *p* < 0.05, 4DUS vs. LSFM, paired *t*-test (*p* = 0.044, IIIR; 0.041, IVR; 0.035, VIR; 0.045, VIL). ‡ *p* < 0.05, LSFM vs. nanoCT, Welch’s *t*-test

**Table 5.**
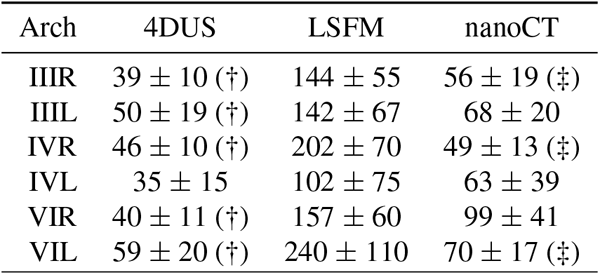
Raw values of spatial-mean peak systolic WSS (in dyne/cm^2^) across each arch based on each imaging modality. All data are presented as population mean ± standard deviation (*n* = 5). † *p* < 0.05, 4DUS vs. LSFM, paired *t*-test (*p* = 0.015, IIIR; 0.0085, IVR; 0.011, VIR; 0.029, IIIL; 0.019, VIL). ‡ *p* < 0.05, LSFM vs. nanoCT, Welch’s *t*-test (*p* = 0.031, IIIR; 0.012, IVR; 0.035, VIL)

We examined the evolution of pressure and WSS at peak flow along each arch at the same 11 evenly-spaced cross-sections as our morphological analyses. We normalized cross-sectional pressure and WSS (spatially average values normalized to arch mean) to highlight the patterning of the forces (Fig. 5C-D). These relative metrics are valuable in identifying regions with particularly elevated or reduced hemodynamic stresses, highlighting potential sites of major developmental events. Normalized values revealed remarkable agreement between relative pressure distributions recovered from nanoCT and LSFM, with no significant difference found at any section along the arches (Table 3, Supplemental figure S6). 4DUS-based simulations featured noticeably smaller pressure drops across all arches with a large number of sections showing significant differences in relative pressure value (Table 3, Supplemental figure S6). Normalized WSS levels followed similar trends across all three modalities with only a few sections showing statistically significant differences (Fig. 5D, Table 3, Supplemental figure S7).

## Discussion

The use of quantitative imaging and computational modeling in development and the biomedical sciences is growing as researchers strive to complement their experimental findings with mathematical calculations to estimate difficult to measure values^17,32,33^. Here, we compare popular small animal imaging modalities: high resolution ultrasound, light-sheet microscopy and nano computed tomography, to assess their ability to accurately capture intricate vessel morphologies and yield accurate computational models.

In our previous study, we identified three subsets of HH26 PAA development^26^, HH26 ‘fast’, HH26, and HH26 ‘slow’ that reflected different timing in arch artery vessel evolution, including lengthening rotation and shape changes. HH26-slow arches were notably shorter than HH26 and HH26-fast embryos, and possessed perfectly circular CSAs. As the embryo progressed to HH26 and HH26-fast, CSAs elongated in the vertical (cranial-caudal) direction, with HH26-fast embryos exhibiting more elliptical (particularly in the latter half) intricate arches with smaller CSAs when compared with the HH26 subset. These type of CSA shape changes could not have been observed through 4DUS which systematically inflated and rounded out CSAs. We did not divide embryos into subsets for this study to maintain focus on modality comparisons.

An advantage of this study is that the exact same embryos were used for LSFM and 4DUS allowing for a direct comparison between these imaging modalities. In addition to inflated CSAs, 4DUS anatomical models and CFD results featured changes in OFT rotation angle and PAA flow split. Kowalski et al^31^ showed that the orientation of the OFT is critically important to flow distributions among the arches. Unsurprisingly, nanoCT embryos, whose cohort featured smaller OFT rotation angles, possessed vastly different arch flow splits from that of the LSFM and 4DUS cohort. Despite cohort-based embryo differences in size and vessel maturity, LSFM embryos were still able to capture the delicate morphology of the arches. Normalized LSFM- and nanoCT-based cross-sectional area, ellipticity, WSS, and pressure results showed strong agreement, while 4DUS differed in all metrics except WSS. This suggests that LSFM faithfully and reliably reproduces the known physiological state of avian PAAs at HH26. The capability of LSFM to reliably capture complex tortuous vessels can be extended to differently-staged avian embryos, embryos of other species, and other small animal samples.

4DUS, in its current form, is not a comparable tool for producing 3D anatomical models of small tortuous vascular geometries as shown here with the large discrepencies captured between LSFM and 4DUS imaging of the same embryo. Inaccuracies in vessel shape and pressure distribution measurements may obscure subtle morphological features that are important for understanding disease propagation. 4DUS has already been proven to be a valuable tool in capturing cardiac chamber anatomy and functionality as well as relative changes in volumes/mass between wild-type and treatment groups, or over the cardiac cycle within the same animal model^22,25^. A unique advantage of 4DUS over LSFM and nanoCT is its *in vivo* live imaging capabilities. It is the most widely accessible and economic imaging modality capable of resolving motions of the heart and outflow tract both within a cardiac cycle^20,34,35^ and between developmental stages^36^. 4DUS provides key cardiac chamber functional metrics and enables wall-motion-informed computational simulations. Future developments, such as advancements in ultrasound technology and more sophisticated post-processing algorithms, can further improve the accuracy and reliability of 4DUS in small animal vascular quantitation.

While both nanoCT and LSFM have shown promise in capturing the intricate morphological details of the HH26 PAAs with great resolution, they are not without limitations (Table 2). Both modalities are limited in their field of view. Imaging of larger vascular systems may require tiling or use of larger voxel size machines (such as µ-CT). Clearing of samples containing mineralized tissues including bone is also challenging. While the sample preparation and imaging techniques used here can be used for any small animal model, the three imaging modalities were tested on the intricate PAA system of the chick embryo animal model. Although the PAAs offered a wide range of diameters, areas and curvature, a limitation of the study is that the vessel wall of the PAAs is not fully developed and does not have organized smooth muscle cell layers^37,38^ at the HH26 stage which may have affected 4DUS image acquisition. The 4DUS live imaging of the pressurized vascular system may have inflated the vessels due to the soft vessel wall composition at timepoint used. The PAAs presents particular challenges for 4DUS imaging as they are surrounded by dense solid mesenchyme, leading to reduced contrast and higher background noise^20^. In neonate and adult animal models, the wall structure, connective tissue compositions, vessel diameter, and curvature may play a role in the quality of 4DUS images. The possibility of simultaneous B-mode ultrasound and ECG recording for adult model animals enables more precise gating and synchronization, though there may also be more sample noise with the more developed tissues^24^.

In summary, LSFM is able to capture high-resolution images for small animal vascular quantitation beginning at the embryonic stage. 4DUS looses some vessel shape and cross sectional area features along complex vessels. More neonate and adult vascular comparison studies will be needed to fully elucidate the limits of 4DUS 3D vascular quantitation. The accuracy of acquired 3D anatomical reconstructions is of critical importance to the integrity of subsequent computational modeling. Given the observed differences in morphology and simulation derived hemodynamic forces between 4DUS, LSFM, and nanoCT, we encourage investigators to use LSFM or computed-tomography for 3D anatomical model reconstructions of vessels. We recommend 4DUS continue to be used for wall motion capture and relative volume changes and suggest that 4DUS-based anatomical models with rigid walls be extensively validated for blood vessels when looking to obtain empirical values.

## Methods

### Embryo Preparation

Fertilized chicken eggs were obtained from commercial farms and incubated in a continuous rocking incubator at 37.5°C, 55% humidity to the desired Hamburger Hamilton stage, HH26 (day 5). Chicks undergoing 4DUS were open-cultered at HH18 (day 3)^39^ and returned to the incubator to continue developing until the desired time point.

### 4D Ultrasound Image Acquisition

4D ((*x, y, z*) and *t*) recordings of chick embryo pharyngeal arch arteries were capturing through a series of B-mode 2D cine-loop recordings ((*x, y*) and *t*) at a multiple imaging planes (*z*). Imaging was performed with a Vevo F2 preclinical ultrasound (FUJIFILM Visualsonics, Toronto, Ontario, Canada) equipped with the UHF71x transducer (Fig.6). The transducer was mounted vertically, perpendicular to the surface of the embryo. A thin layer of Tyrode’s Solution was added between the embryo and the transducer to act as an aqueous conduit. To maintain warmth during scan sessions, the base of the cup containing the embryo was mounted in a waterbath operating between 40 and 42 ºC. The waterbath was placed atop a microadjustable stage that allowed for embryo translation. The direction of translation (*z*) was aligned along the dorsal-ventral axis of the PAAs (Fig. 1B) so that the axial cross-sections of the vessels were captured by each image plane. The imaging window was adjusted to maintain a in-plane resolution of 7.3 to 8.7 µm and with 50 µm z-steps along the dorsal-ventral axis. Each B-mode cine-loop recording was acquired at a frame-rate of 200-300 frames per second for at least 5000 frames to capture a minimum of 20 cardiac cycles.

**Figure 6.**
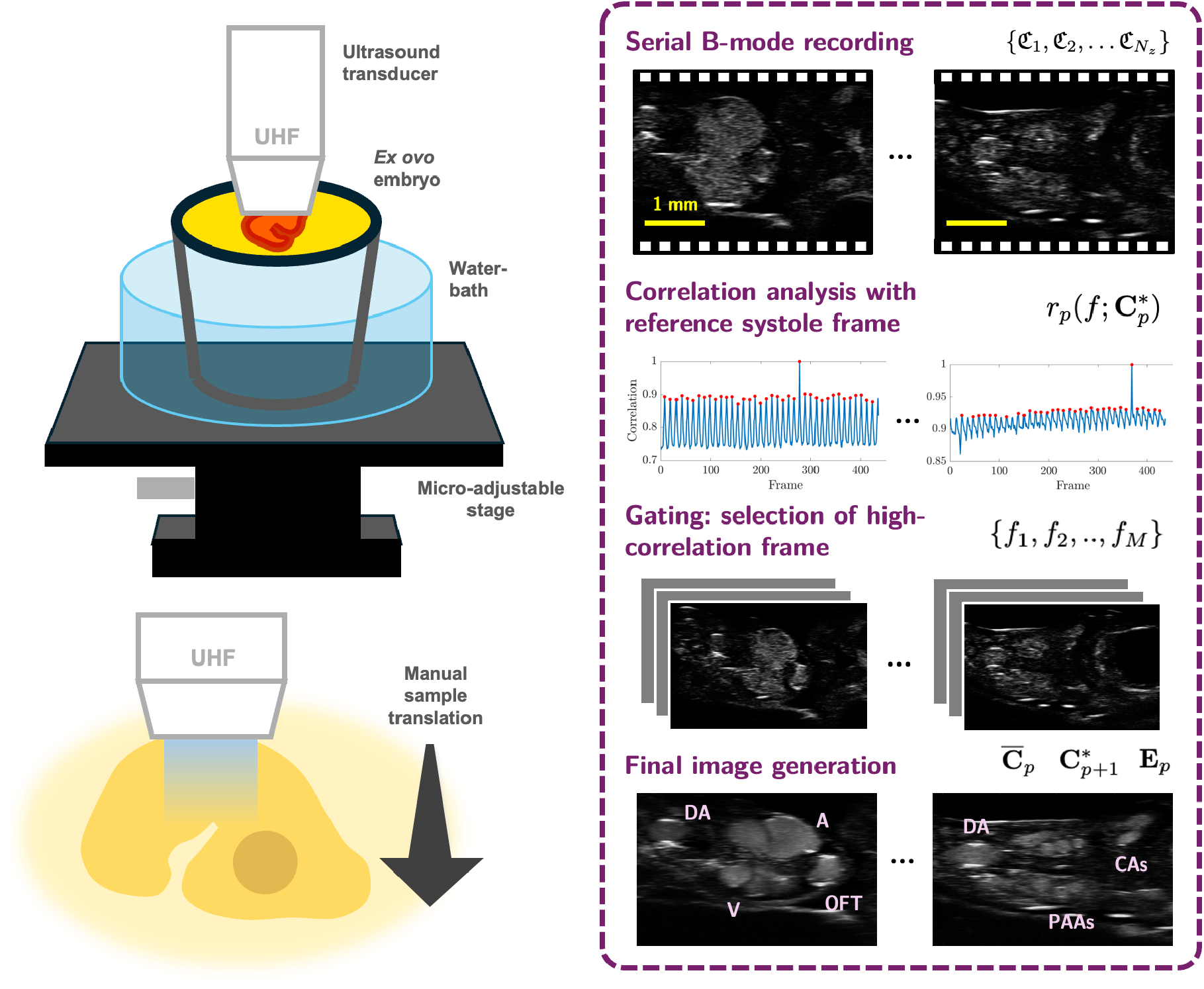
4DUS imaging setup and retrospective gating. The *ex ovo* live embryo is placed in a miniature waterbath, and the ultra high-frequency ultrasound transducer is fixed perpendicular to the embryo body. A microadjustable stage is used to translate embryo in relation to the transducer to select imaging planes. Gating of a cine loop (right panel) at a given imaging plane, and synchronization of two consecutive cine loops, is done through correlation analysis with a reference frame. High correlation frames are used to construct a still imaging of the embryo at the cardiac phase of interest with enhanced blood-space pixel intensity as shown in the final image generation.

Due to the difficulty in simultaneously imaging and recording avian ECG, a 4DUS processing method without ECG gating was adopted from^20^ (Fig.6). Briefly, a total of *N*_*z*_ cine-loop recordings were recorded along the (*z*) axis with each recording being *N*_*x*_ × *N*_*y*_ × *N*_*t*_ (width × height ×number of frames). From these recordings, a compiled 4DUS dataset or collection of arrays 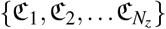 was generated with ℭ_*p*_ being a *N*_*x*_ × *N*_*y*_ × *N*_*t*_ three-dimensional array representing the cine-loop recording at the *p*-th imaging plane. Frame *f* of this collection is designated as 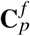 with the intensity of the (*i, j*) pixel included as 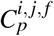. From this dataset, we calculated the average 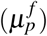 and standard deviation 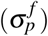 of pixel intensities of a frame *f* from collected at imaging plane *p*.

A retroactive gating and synchronization algorithm with frame-to-frame correlation analysis was used to enhance vascular region contrast. As part of the gating process, a reference frame 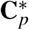 was defined at a peak systole for each imaging plane *p*. The reference frame for the first imaging plane 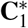 was manually selected based on direct inspection. The similarity between all frames in ℭ_*p*_ and the reference frame was assessed with a correlation metric *r*_*p*_ defined as

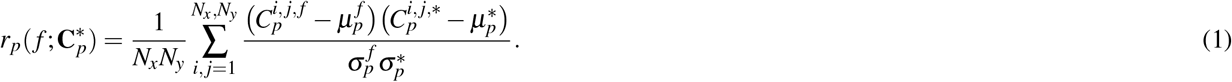

Assuming that embryo motion during recording was caused only by periodic heart motion, *r*_*p*_ was expected to be periodic in *f*. We invoked a peak-finding algorithm to find a set of *M* frame numbers *ℱ* = { *f*_1_, *f*_2_, .., *f*_*M*_} at which sufficiently regular and prominent local maxima of *r*_*p*_ occur. This list identified the frames most similar to the reference frame, and most likely to represent peak systole. We then calculated the mean peak-systole frame 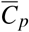 with

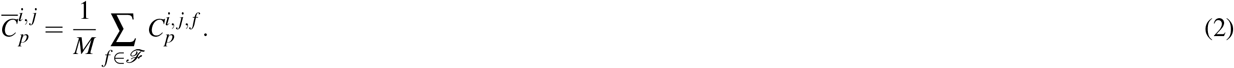

The reference frame for the next imaging plane 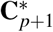 was chosen to be the frame that maximizes 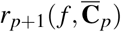, i.e. showed the greatest similarity to the previous mean peak-systole.

We leveraged the fact that B-mode ultrasound recording registers blood space with more dynamically fluctuating pixels than solid tissue to construct a still image with enhanced blood space pixel brightness and contrast peak systole (**E**_*p*_) at plane *p* (Eqn 3).

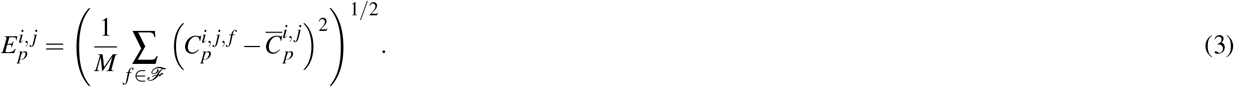

In the event that the blood space did not register with a noticeably higher pixel intensity, we scaled **E**_*p*_ and 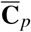 to be the same mean pixel intensity and summed them together. We then stacked 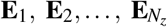 together to form the motion-corrected 3D image of the embryo vasculature at peak systole.

### Lightsheet Microscopy Sample Preparation and Image Acquisition

Light sheet fluoresence microscopy (Fig. 7) was performed on the same embryos used for 4DUS. Embryos were sacrificed within three hours of 4DUS image acquisition and transferred into warm Tyrode’s solution. Using micro-needles fashioned from pulled capillary tubes (0.75 mm ID) cut to 20-35 µm inner diameter via a microforge (Glassworx, St. Louis, MO) and a micromanipulator (model M3301L, World Precision Instruments) the embryo’s vascular system was flushed with warm Tyrode’s solution, endopainted^40^ with FITC-poly-L-lysine and perfusion fixed with 4% (w/v) paraformaldehyde (PFA, Sigma-Aldrich, St Louis, MO, USA) to preserve inner vascular volumetric integrity. The embryo was subsequently dehydrated through a series of methanol steps and optically cleared in Ethyl cinnamate (ECi) as detailed in our endo-iDISCO protocol^18^. LSFM imaging of the embryos was performed using a Zeiss Lightsheet Z.1 system (Carl Zeiss, Oberkochen, Germany). The optically-cleared embryo was affixed to a glass capillary mounted into a sample holder and lowered into an imaging chamber filled ECi. The samples were illuminated with a light-sheet thickness of 5µm and imaged with a 5X objective set to 0.8 zoom. These settings resulted in an in-plane resolution of 1.3µm and an optimal z-step size of 1.9µm. To keep the dataset file size manageable, we imaged the embryos in roughly the sagittal direction (Fig. 1C).

**Figure 7.**
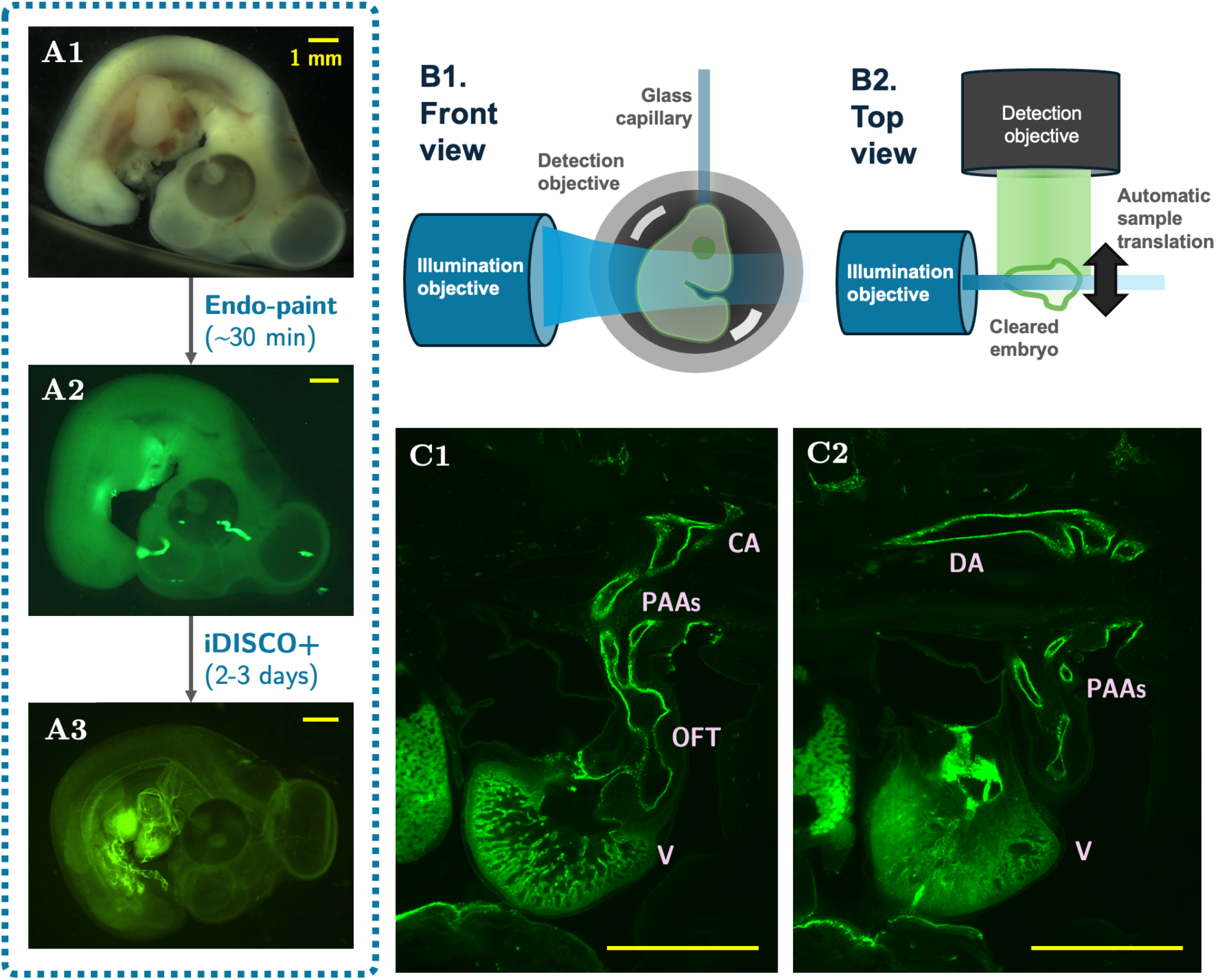
LSFM sample preparation and imaging setup workflow. Widefield images of the embryo (A1) after dissection, (A2) after endo-painting, (A3) after clearing with modified iDISCO+ .(B1) Interior view of the Zeiss Lightsheet Z.1 scope during imaging (B2) Relative position of the embryo to the mounting glass capillary, illumination objectives, and imaging objective. (C1-2) In-plane LSFM images of the pharyngeal region of HH26 embryos. Vessels are labeled with green fluorescence, and vessel names are annotated in pink. Scale bars indicate 1 mm. ECi: ethyl cinnamate, CA: carotid artery, PAAs: pharyngeal arch arteries, OFT: outflow tract, V: ventricle, DA: dorsal aorta

### NanoCT Sample Preparation and Image Acquisition

Embryos and high-resolution images were obtained from our previous 2018 study^26^. Briefly, embryos were perfusion fixed using micro-needles fashioned from pulled capillary tubes (0.75 mm ID) cut to 20-35 µm inner diameter via a microforge (Glassworx, St. Louis, MO). A micromanipulator (model M3301L, World Precision Instruments, Sarasota FL) was used to position the needle into the apex of heart and flush the embryos vascular system with phosphate buffer solution followed by 4% paraformaldehyde to preserve inner vascular volumetric integrity. The embryos were then left in 4% paraformaldehyde for 24-48 hours and stained via a diluted form of Lugol solution, aqueous potassium iodide and iodine, (Sigma-Aldrich, L6146). Embryos were dehydrated down to 100% ethanol placed in polymerase chain reaction tubes and sent to undergo 3-4µm nano-computed tomography scans (nano-CT) via Zeiss-Xradia Versa 520 X-ray microscope.

### Zero-dimensional model

Zero-dimensional (0D) electric analog representations of the PAA system were created for each modality-based cohort embryo to facilitate tuning of boundary conditions. Lumped parameter models were simulated using the SimVascular svZeroDSolver^41^ with Windkessel (RCR) outflow boundary conditions. The full 0D model consisted of the arch artery system, including the outflow tract (OFT) inlet, septatated portion of OFTs, three arch pairs, cranial and caudal sections of the dorsal aorta (DoA), and the distal components associated with these outlets (Fig. 3).

### In-silico Model Preparation and Morphological Analysis

Both LSFM image stacks and 3D vascular images extracted from 4DUS recordings were converted to .vti image files using ImageJ and Paraview. Special precaution was taken to encode the voxel size to reflect the difference between in-plane and axial resolution for both imaging modalities. The images were imported to the open-source cardiovascular modeling software SimVascular where the PAA manifolds were constructed through lofted 2D segmentation^42^. PAA models derived from nanoCT were reconstructed via MIMICS (Materialise), 3MATICs (Materialise) and Geomagics Studio 10 (Geomagic) as described in^26^.

NanoCT and LSFM PAAs were scaled to account for the difference between dehydrated and native vessel size, as determined by a series of in vivo diameter verifications. India ink and Texas Red Dextran were used to obtain native vessel size, results were compared with that of 3D reconstructions and a scaling factor for each modality was generated. Additionally, LSFM PAA manifolds were directly compared to 4DUS PAAs as the same embryos were used for live 4DUS and cleared LSFM. The Visualization Toolkit (VTK) and the Vascular Modeling Toolkit (VMTK)^43^ were used for morphological analysis including computing of centerlines, ellipticity and vessel cross sections along the centerlines.

### Multiscale Blood Flow Simulation Procedure

We simulated hemodynamics in the five subject-specific models derived from each imaging modality by solving the three-dimensional Navier-Stokes equations using SimVascular’s stabilized finite-element solver. Blood was treated as an incompressible Newtonian fluid with a density of *ρ* = 1060 kg m^−3^ and a kinematic viscosity of *µ* = 3.71 cP. Vessel walls were assumed to be rigid and impermeable with no-slip boundary conditions. Models were meshed using TetGen to have a tetrahedral mesh with approximately 120,000 mesh points (around 660,000 elements), following a mesh convergence analysis. A Doppler velocity based pulsatile flow inlet waveform, taken from our previous study^26^ was used for the nanoCT embryos and a newly acquired Doppler-based flow curve used for 4DUS and LSFM cohort (Fig S2) embryos. A plug inlet flow profile was assumed. Following a first round of 3D blood flow simulations, subject-specific lumped parameter (0D) circuit models were used to tune the 3D-0D models to obtain 90-10 caudal-cranial flow splits over the course of each cardiac cycle. DoA pressure was tuned to match published values^44^. Multiple iterations of updating the lumped parameter model, calculating new Windkessel bounds, re-running the 3D-0D simulation with the new bounds and updating the resistance values of the lumped parameter model were required to tune all models (Fig. 3).

## Ethics approval

Human Ethics and Consent to Participate declarations: not applicable. Institutional Animal Use and Care Committee, IACUC : not applicaable.

## Availability of data and materials

The data that supports the findings of this study are available within the article and its supplementary material. Computational models built for this study will be made available upon request to the corresponding author.

## Code availability

The computational fluid dynamics software used, SimVascular, is open source: www.simvascular.org. Inquiries regarding the custom 4DUS post-processing and PAA vascular property analysis pipeline should be directed to the corresponding author, SEL.

## Acknowledgements

The authors would like to thank Dr. Bobby Thompson for his introduction to endo-painting and Ritika Singh for early light-sheet microscopy work. We thank Dr. Marcella Erb and Jennifer Santini of the UCSD School of Medicine Microscopy Core (supported by NINDS P30NS047101) for their assistance in LSFM sample preparation and imaging. The authors also thank Prof. Jesse Jokerst and group members for managing the ultrasound scanner (funded by grant S10 OD032268).

## Author contributions statement

Conceptualization (SEL), Funding acquisition (SEL), Methodology (SEL, DZ), Supervision (SEL) Writing - review & editing (SEL, DZ), Data curation (DZ), Formal analysis (DZ), Investigation (DZ), Writing - original draft (SEL, DZ).

---

**Figure S1:**
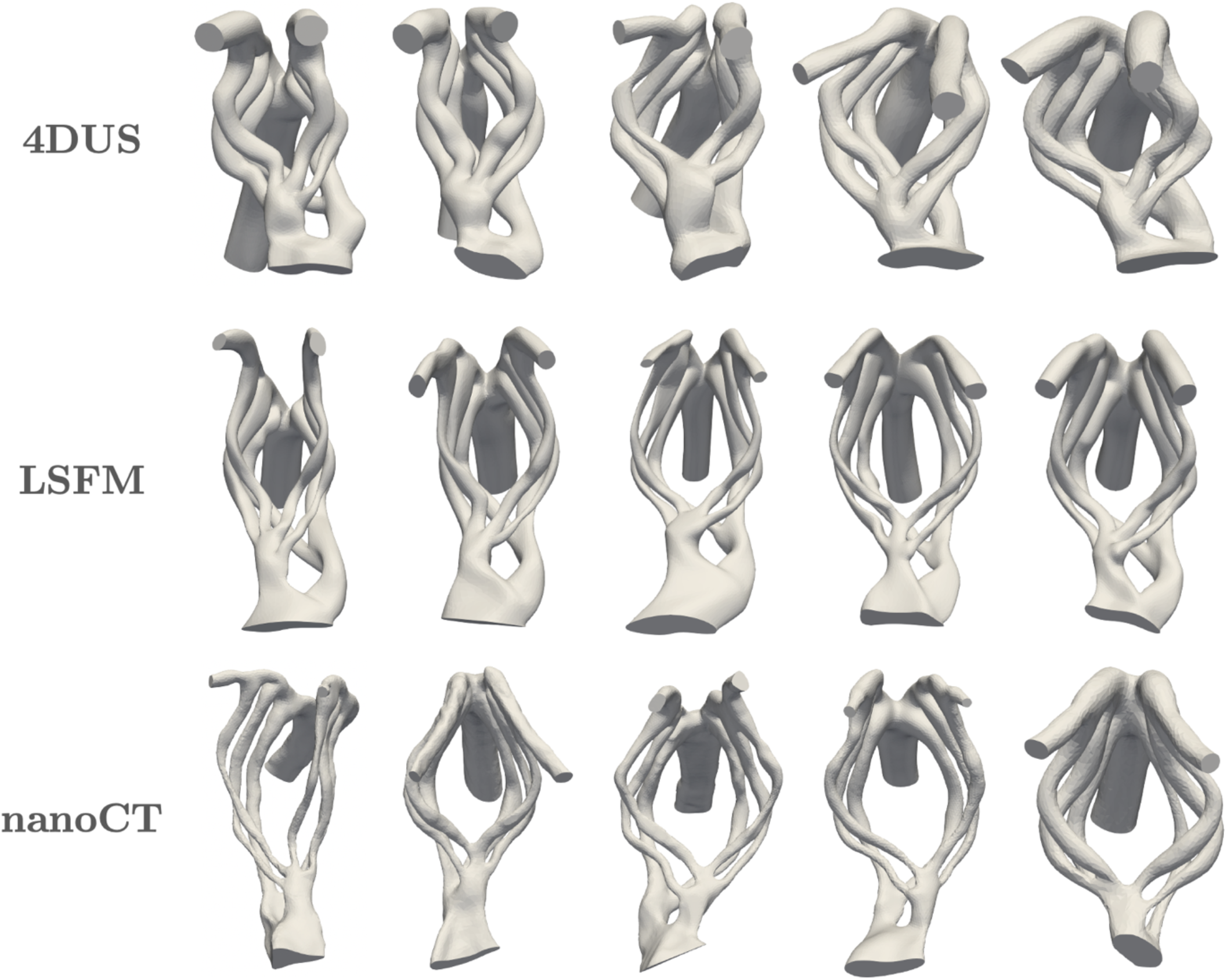
Subject specific anatomical reconstructions of control HH26 chick embryo pharyngeal arch arteries (PAAs) obtained from four-dimensional ultrasound (4DUS), light sheet fluorescence microscopy (LSFM), and nano computed tomography (nanoCT).

**Figure S2:**
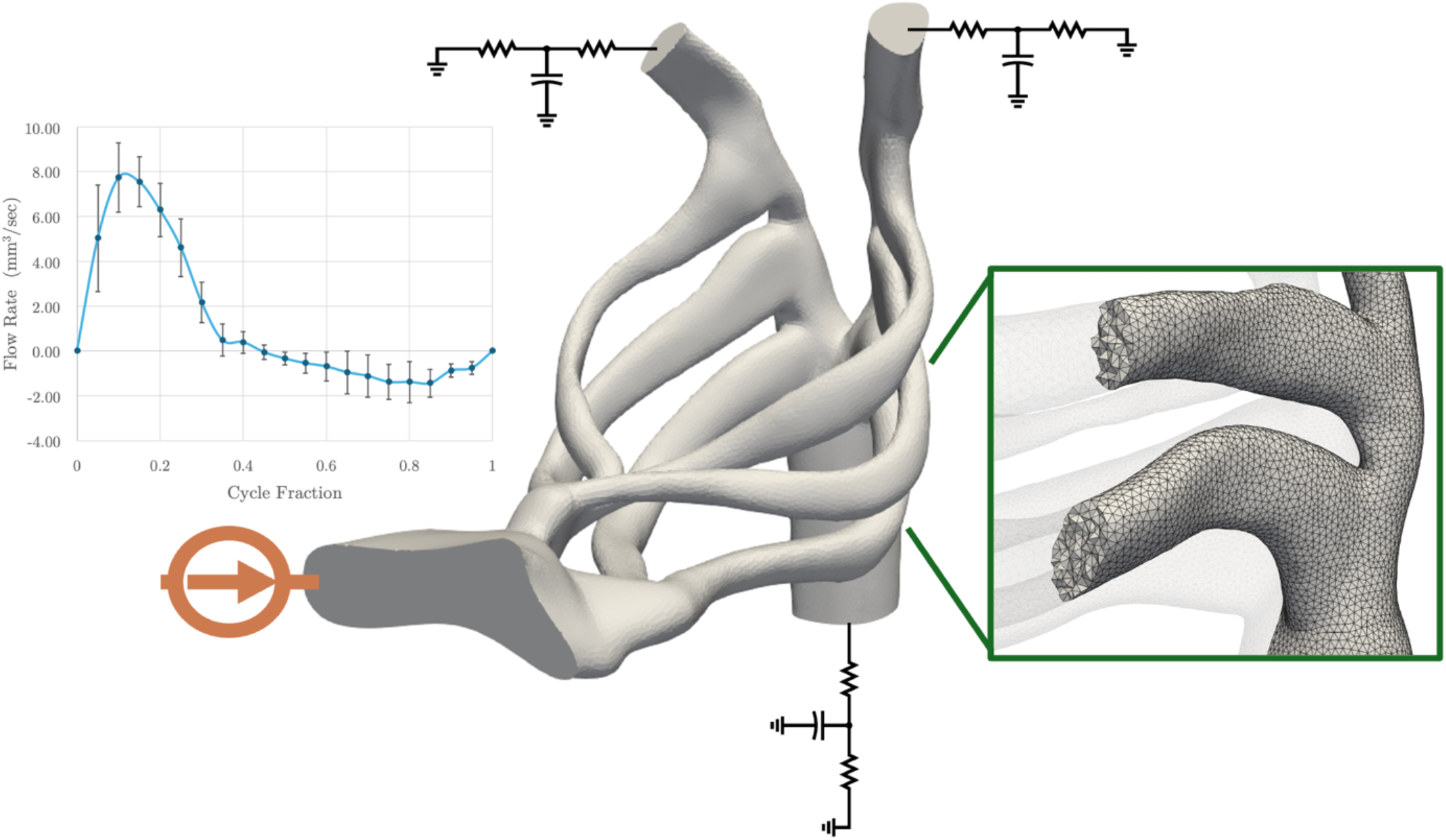
Model preparation for computational fluid dynamic simulations of pulsatile blood flow in HH26 PAAs. The anatomical reconstruction models are discretized into tetrahedral mesh. A flow curve derived from Doppler ultrasound measurements of control HH26 cardiac outflow velocities is used as inflow boundary condition. Three-element RCR Windkessel boundary conditions are imposed at the outlets.

**Figure S3:**
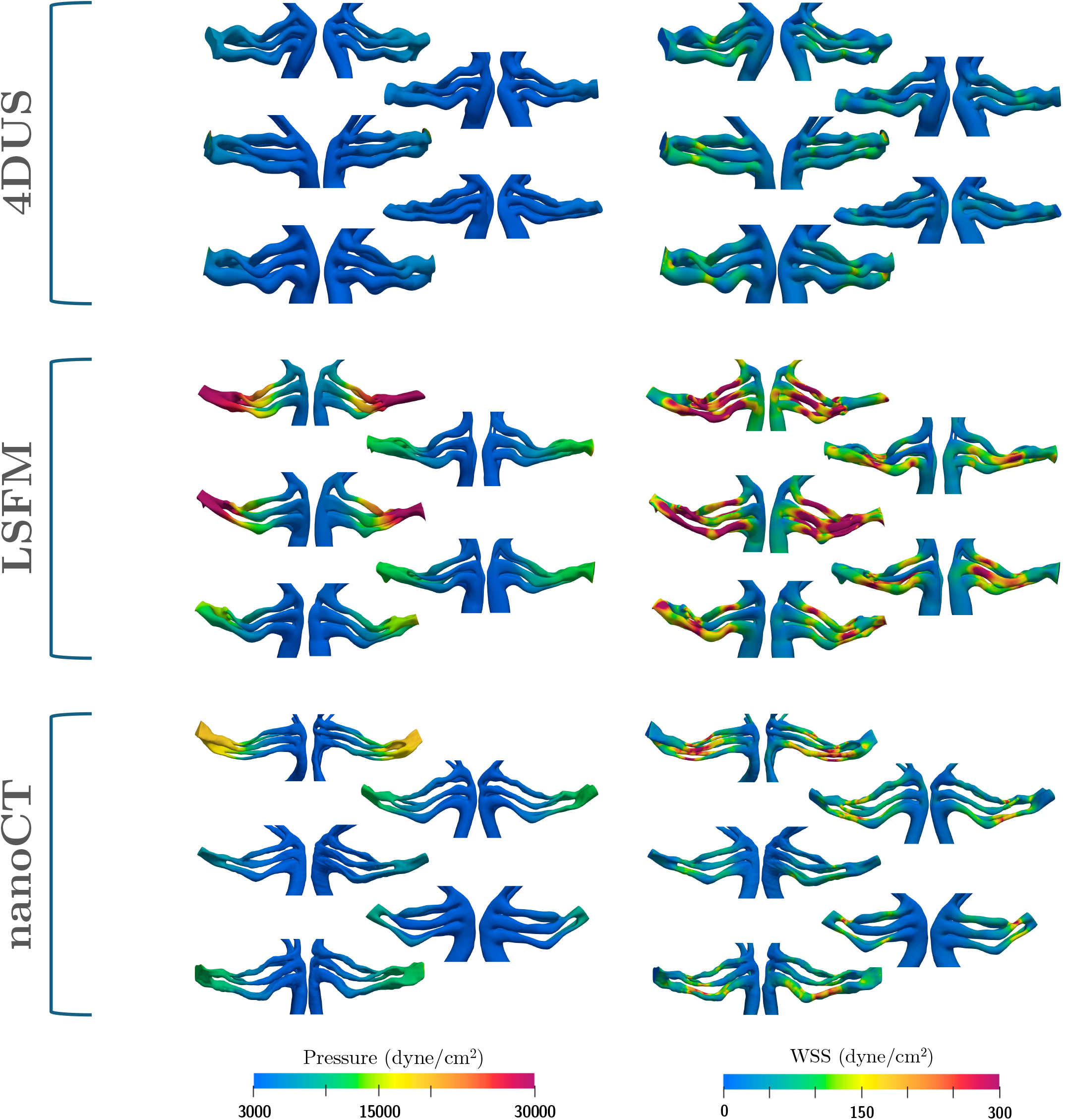
Peak flow pressure and wall shear stress (WSS) distributions of HH26 embryos uncovered by subject-specific hemodynamic simulations

**Figure S4:**
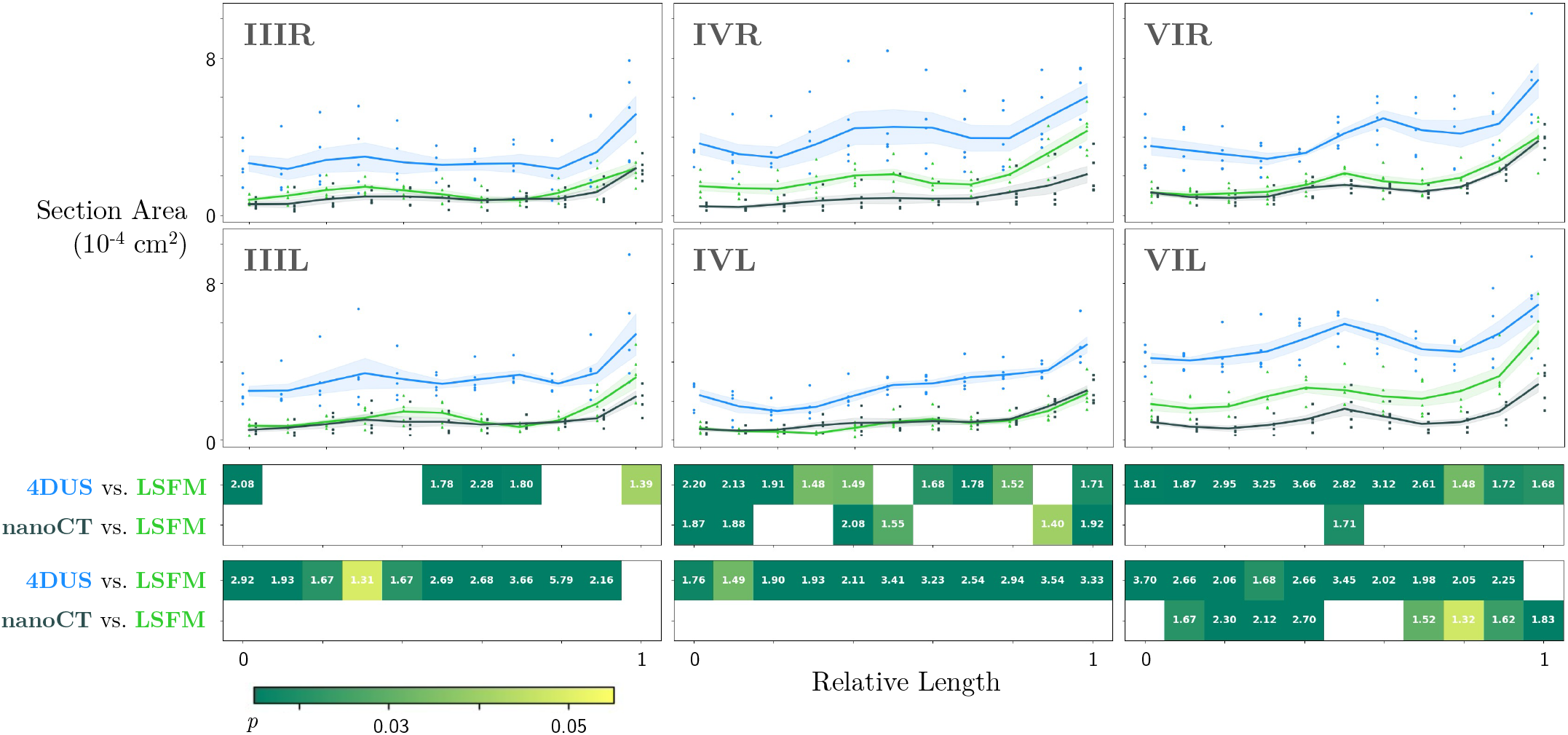
Evolution of cross-sectional areas along each arch, scaled against arch length. Average values of *n* = 5 replicate embryos for each imaging modality are plotted as a solid line, with shaded areas indicating mean ± standard error. Individual datapoints are included as dots. The heatmap indicates locations with statistically significant (*p* < 0.05) differences in cross-sectional areas between the 4DUS and LSFM cohort (paired *t*-test) or the nanoCT and LSFM cohort (Welch’s *t*-test). Annotations indicate -log_10_(*p*-value).

**Figure S5:**
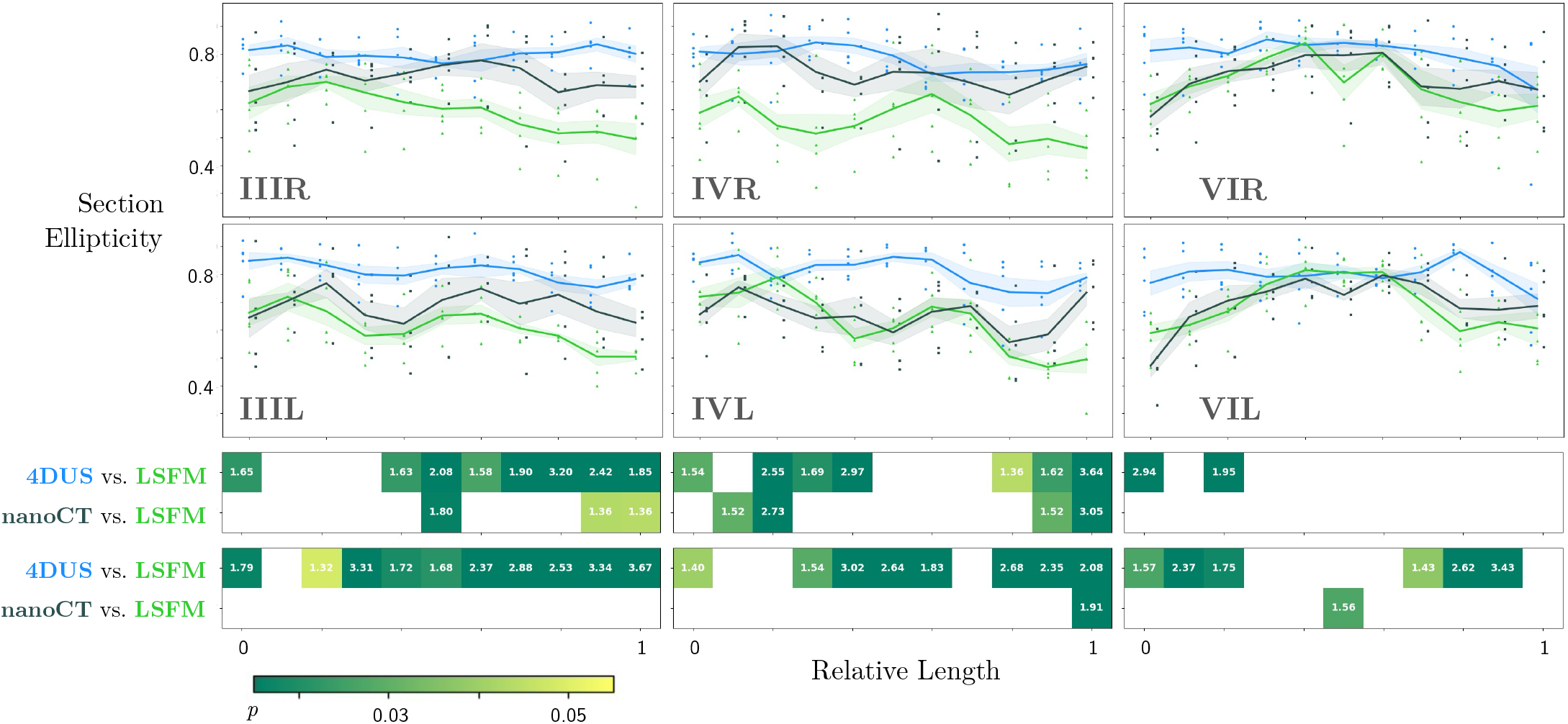
Evolution of cross-sectional ellipticity along each arch. Average values of *n* = 5 replicate embryos for each imaging modality are plotted as a solid line, with shaded areas indicating mean ± standard error. Individual datapoints are included as dots. The heatmap indicates locations with statistically significant (*p* < 0.05) differences in cross-sectional ellipticity between the 4DUS and LSFM cohort (paired *t*-test) or the nanoCT and LSFM cohort (Welch’s *t*-test). Annotations indicate -log_10_(*p*-value).

**Figure S6:**
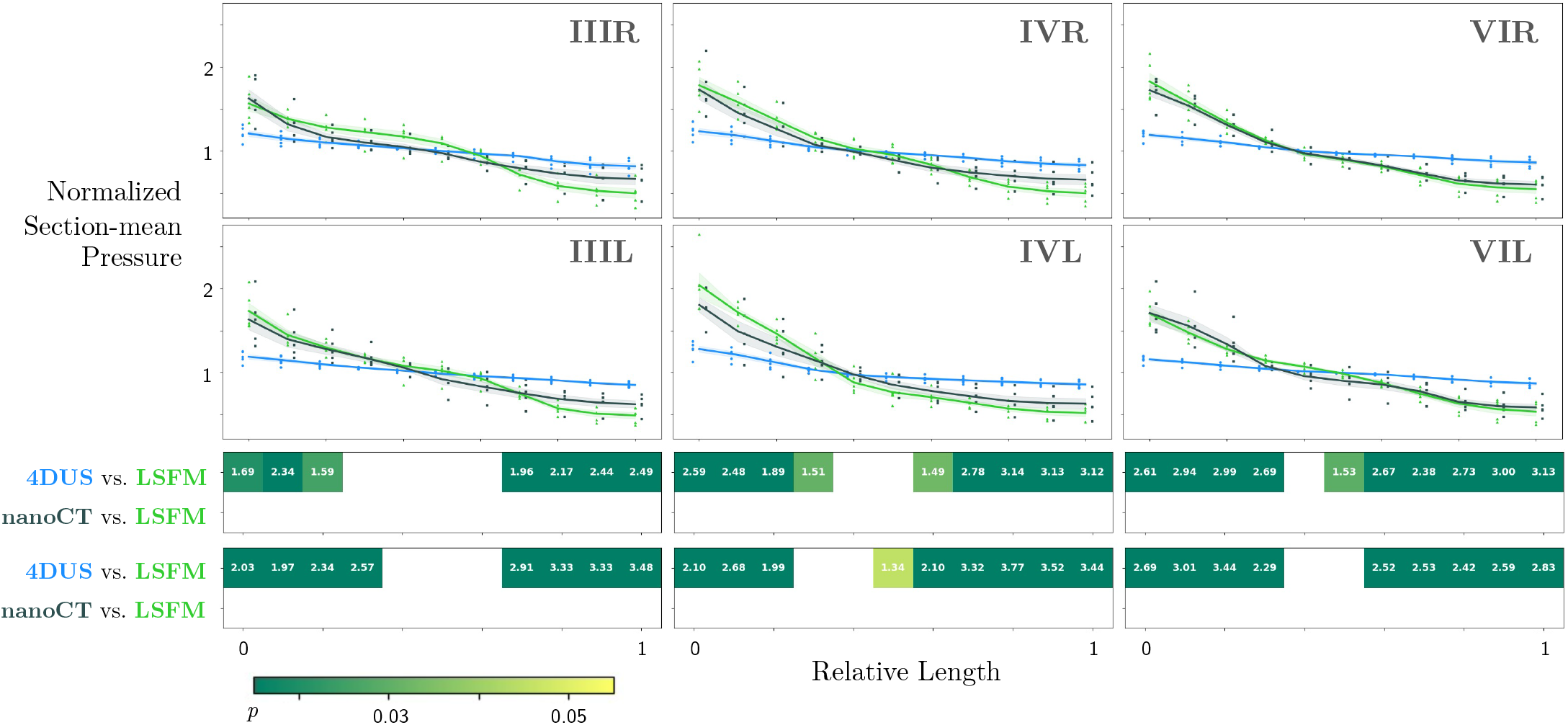
Evolution of cross-section average peak flow pressure along each arch, normalized against the arch mean value for each replicated. Average values of *n* = 5 replicate embryos for each imaging modality are plotted as a solid line, with shaded areas indicating mean ± standard error. Individual datapoints are included as dots. The heatmap indicates locations with statistically significant (*p* < 0.05) differences in cross-sectional normalized pressure between the 4DUS and LSFM cohort (paired *t*-test) or the nanoCT and LSFM cohort (Welch’s *t*-test). Annotations indicate -log_10_(*p*-value).

**Figure S7:**
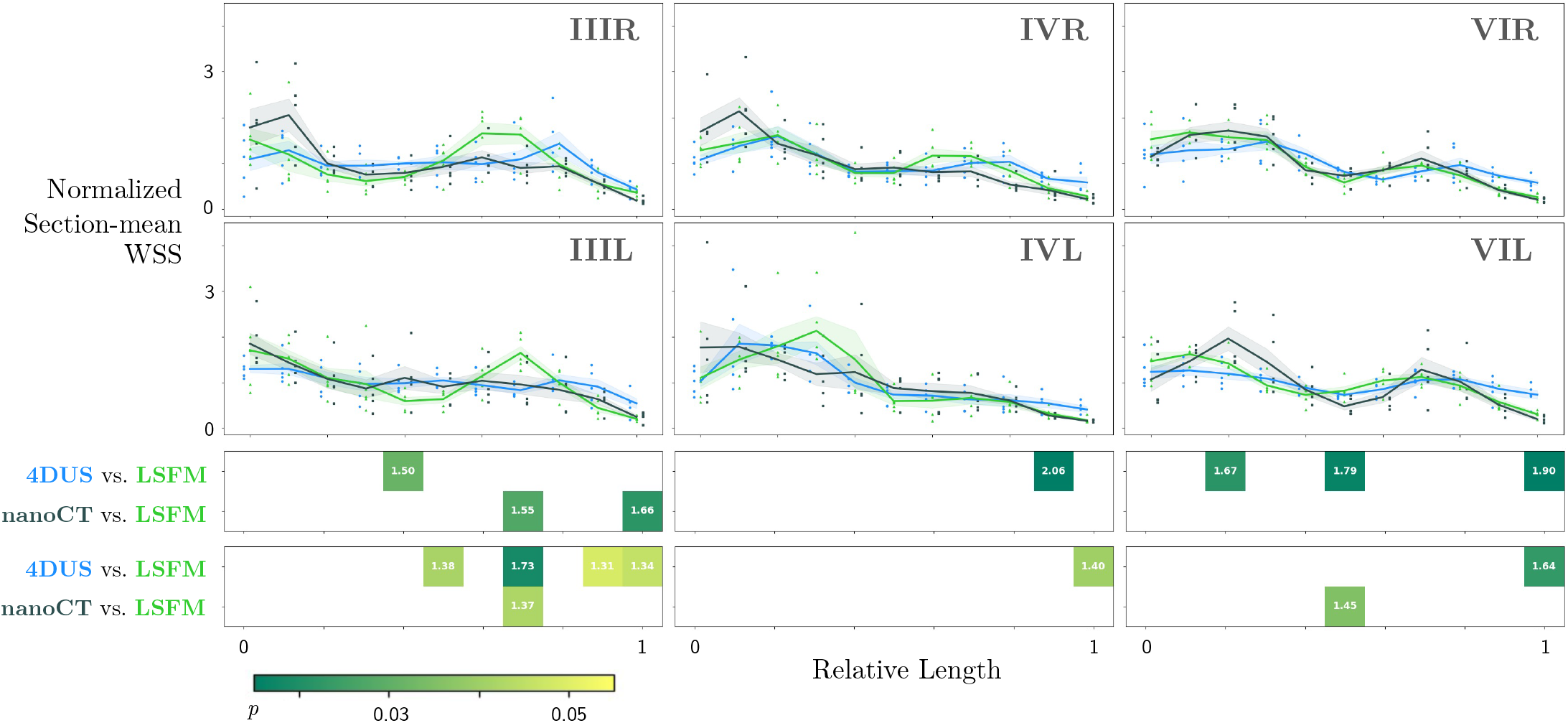
Evolution of cross-section average peak flow wall shear stress (WSS) along each arch, normalized against the arch mean value for each replicated. Average values of *n* = 5 replicate embryos for each imaging modality are plotted as a solid line, with shaded areas indicating mean ± standard error. Individual datapoints are included as dots. The heatmap indicates locations with statistically significant (*p* < 0.05) differences in cross-sectional normalized WSS between the 4DUS and LSFM cohort (paired *t*-test) or the nanoCT and LSFM cohort (Welch’s *t*-test). Annotations indicate -log_10_(*p*-value).

